# Multi-Scale Geometric Network Analysis Identifies Melanoma Immunotherapy Response Gene Modules

**DOI:** 10.1101/2023.11.21.568144

**Authors:** Kevin A. Murgas, Rena Elkin, Nadeem Riaz, Emil Saucan, Joseph O. Deasy, Allen R. Tannenbaum

## Abstract

Melanoma response to immune-modulating therapy remains incompletely characterized at the molecular level. In this study, we assess melanoma immunotherapy response using a multi-scale network approach to identify gene modules with coordinated gene expression in response to treatment. Using gene expression data of melanoma before and after treatment with nivolumab, we modeled gene expression changes in a correlation network and measured a key network geometric property, dynamic Ollivier-Ricci curvature, to distinguish critical edges within the network and reveal multi-scale treatment-response gene communities. Analysis identified six distinct gene modules corresponding to sets of genes interacting in response to immunotherapy. One module alone, overlapping with the nuclear factor kappa-B pathway (NFKB), was associated with improved patient survival and a positive clinical response to immunotherapy. This analysis demonstrates the usefulness of dynamic Ollivier-Ricci curvature as a general method for identifying information-sharing gene modules in cancer.

## Introduction

Melanoma is an aggressive cancer which arises due to dysregulation of melanocytes, typically in the skin^1^. This dysregulation generally involves mutational, epigenetic and transcriptional changes that affect multiple proteins and molecular subsystems in the cancer cell, driving tumorigenesis and growth^2-6^. Although numerous studies have examined the expression profiles of melanoma, the mechanisms of therapeutic response and resistance are not thoroughly characterized^7-9^. In this study, we focus on transcriptomic responses to the immunotherapy drug nivolumab, a monoclonal antibody that functions via anti-PD1 immune checkpoint blockade^10^.

Given the complex interplay of molecular components involved in the therapeutic response to immunotherapy in cancer cells, network models offer a method to study cancer from a systems perspective. In a network model, individual genes and pairwise relationships between genes can be represented as vertices and edges, respectively. Previous studies have demonstrated that network geometric properties including network curvature and entropy can, for example, distinguish melanoma cells from normal cells, suggesting network geometry may be useful to classify cancer and non-cancer cells and investigate molecular changes that occur during tumorigenesis or in response to treatment^11-14^.

By considering correlations among experimentally-measured gene expression data, one can construct a weighted network that can identify key modules of genes with correlated expression patterns that may be implicated in cancer therapeutic response^15, 16^. Such gene modules might indicate coordinated transcriptomic programs related to certain cellular responses, for example cancer signaling or metabolic pathways that may contribute to treatment success or failure. Commonly, gene correlation network analyses involve potentially arbitrary cutoffs of inter-gene correlation values or, alternatively, principal component analysis to identify such gene modules^15, 17^. However, few methods incorporate distance information on the network in a natural way. One method that does this is Ollivier-Ricci curvature, a network geometric measure of the connectedness of neighborhoods, which has been shown to distinguish within-cluster (positive) and between-cluster (negative) connections, and thereby can be used to define interconnected gene modules in a robust way^18-20^. In fact, previous research utilized such network geometric approach to study pediatric sarcoma and identify novel functional gene associations including the EWSR1-FLI1-ETV6^21^.

In this study, we applied a multi-scale geometric network analysis approach to study the transcriptomic response of melanoma tumors treated with the immunotherapy drug nivolumab. We modeled changes in gene expression as a weighted correlation network and assessed network geometry at multiple diffused scales in order to determine which network edges (correlations) correspond to ‘critical’ connectivity bridges of the network. This allowed us to identify several distinct gene modules, which we subsequently assess in terms of differential regulation in response to treatment and consider clinical associations including mutational subtype and overall survival. Using this approach, we compared pathway analysis of the gene modules to known mechanisms and identified a potential key marker of melanoma immunotherapeutic response.

## Results

### Multi-Scale Network Analysis Identifies Gene Modules in Melanoma Differential Gene Correlation Network

We focus this study on a publicly available melanoma transcriptomic dataset measuring gene expression (by RNA-seq) in melanoma patient tumors before and during treatment with the immunotherapy drug nivolumab, a PD-1 immune checkpoint inhibitor^10^. In 43 patients, gene expression was measured in both pre- and on-treatment conditions, allowing analysis of the matched expression changes (on-treatment minus pre-treatment) in response to therapy for each patient. We considered only genes relevant to immunotherapy response by narrowing our focus to 912 genes, including 18 genes with known involvement in immunotherapy response to PD1 blockade and 894 genes sharing pairwise interactions with those 18 core genes in a known protein-protein interaction database (STRINGdb)^22-25^. To model co-expression relationships among these genes, we constructed a correlation network based on the Pearson correlation of each pair of genes that shared a protein-protein interaction, for a total of 50,518 edges.

We applied a multi-scale geometric assessment of the correlation network to identify, in an unsupervised manner, clusters of correlated gene modules of melanoma immunotherapeutic response. We first applied a diffusion process to simulate diffusion of information across the network on increasing scales according to a pseudotime/scale parameter *τ* (Fig. 1A). This diffusion allowed us to examine the network through a ‘lens’ of various scales to observe multi-scale properties of the network. We then measured Ollivier-Ricci curvature (ORC) *κ*, a key network geometric property that represents the closeness of two neighboring distributions of connected genes in the network. Larger values of *κ* indicate that information is more closely related between the neighborhoods around two genes of a given edge in the network. ORC is thus valuable to identify which gene correlations are likely within-cluster (defined as *κ* positive) and which are likely between-cluster (defined as *κ* negative).

**Figure 1:**
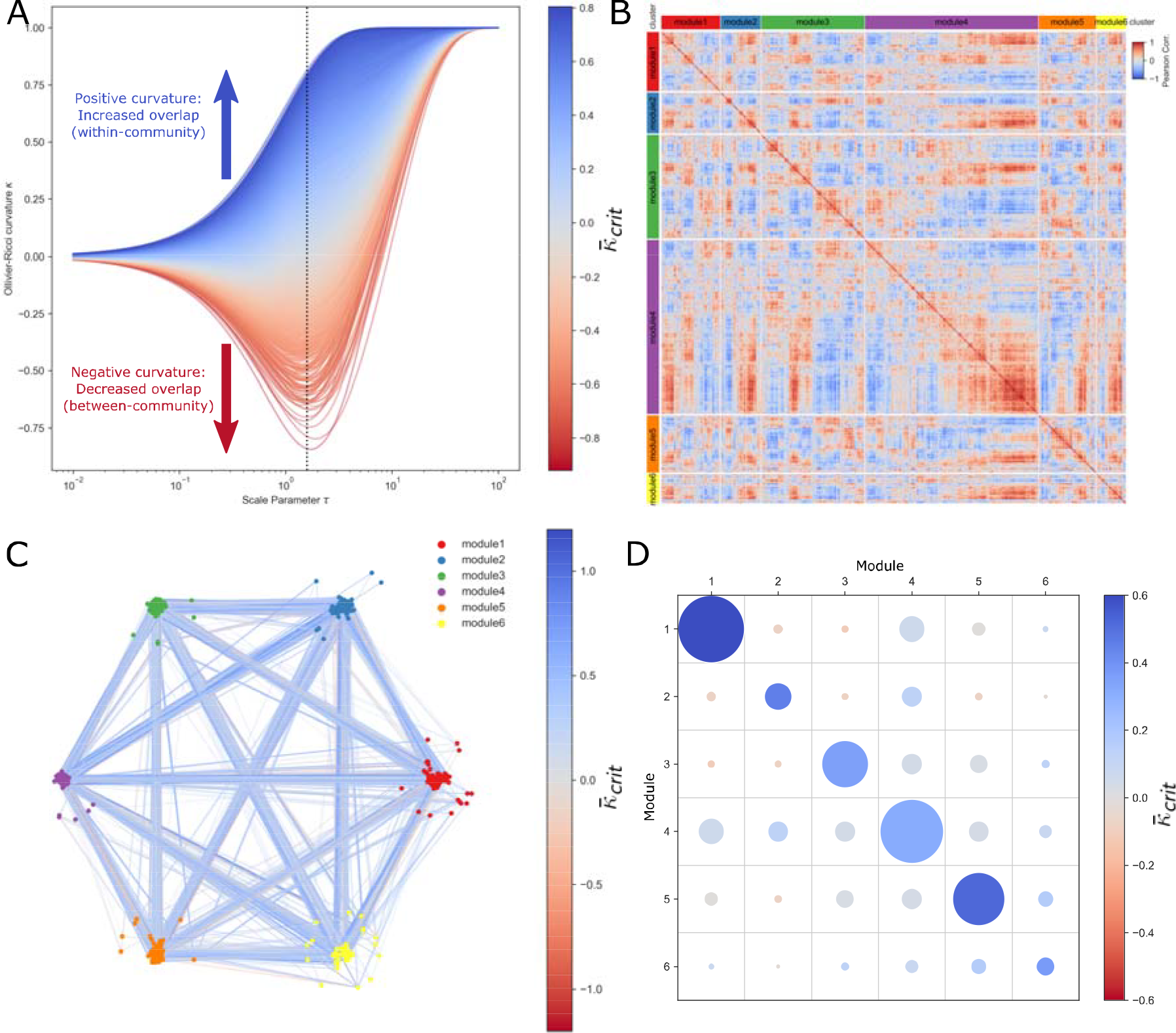
Identification of Correlated Gene Modules in Melanoma Immunotherapy Response with Multi-scale Geometric Network Analysis. **A:** Line plots of gene-gene edge curvature *κ* over diffusion for *τ*, including vertical line critical *τ*_*crit*_=1.58, a point of high discrimination, whereby lines are colored by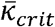. **B:** Correlation heatmap of 6 gene modules identified with weighted Louvain clustering. **C:** Graph network with edges colored by 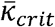 and layout partitioned by cluster. **D:** Average 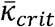 of edges within or crossing between each pair of gene clusters, where color indicates average 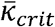 and area of each circle is proportional to number of edges.

Importantly, measuring the curvature of the correlation network at various information diffusion scales allowed us to determine a *τ*_*crit*_ threshold at which to best partition the network into correlated gene modules (Fig. 1A; Table 1). At *τ*_*crit*_ = 1.58, we extracted *κ*_*crit*_ as the curvature of each edge at *τ*_*crit*_ and additionally defined 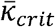 as a ‘smoothed’ estimate of curvature integrated over all diffusion steps up to *τ*_*crit*_. We then applied a weighted Louvain clustering algorithm to partition the network while maximizing average 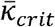(high shared information) within clusters while minimizing average 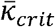(low shared information) between clusters, resulting in six distinct modules of correlated genes (Fig. 1B,C). With the defined clusters, we observed greater 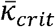 (Fig. 1D; unpaired t-test: p<0.001) among within-cluster edges (n=20,289 edges, mean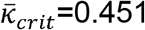) as compared to between-cluster edges (n=4,970 edges, mean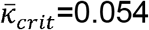), suggesting that ORC can effectively separate correlated gene modules based on network geometry.

**Table 1:** Top five highest and lowest curvature edges at *τ*_*crit*_:

| Gene A | Gene B | $\bar{\kappa}_{crit}$ |
| --- | --- | --- |
| FRK | ARMC8 | 0.804 |
| CDC27 | CBX8 | 0.791 |
| CD3D | CD274 | 0.777 |
| CUL1 | PSMA4 | 0.768 |
| CLTC1 | SYNJ2 | 0.764 |
| LDLR | APOB | -0.921 |
| MAPK10 | TP53 | -0.842 |
| CDKN1A | ABL1 | -0.826 |
| LDLR | APOA1 | -0.822 |
| AP2S1 | SH3GL3 | -0.796 |

Next, in order to highlight relevant biological pathways involving the genes of each module, we performed pathway analysis (Fig. 2A, Supp. Table 1). Gene Ontology (GO) enrichment analysis indicated that each module was associated with distinct pathways (Fig. 2B), which we summarize as follows: Module 1 enriched for endocytosis and vesicle transport. Module 2 enriched for leukocyte chemotaxis and migration. Module 3 enriched for histone modification and chromatin remodeling. Module 4 enriched for cell adhesion and leukocyte proliferation. Module 5 enriched for proteasomal catabolism and ubiquitination. Module 6 enriched for nuclear factor kappa-B (NF-kB) signaling, transcription factor activity, and cytokine production and signaling. These gene modules thus represent distinct biological processes that may be fundamentally modulated by melanoma tumors in response to immunotherapy.

**Figure 2:**
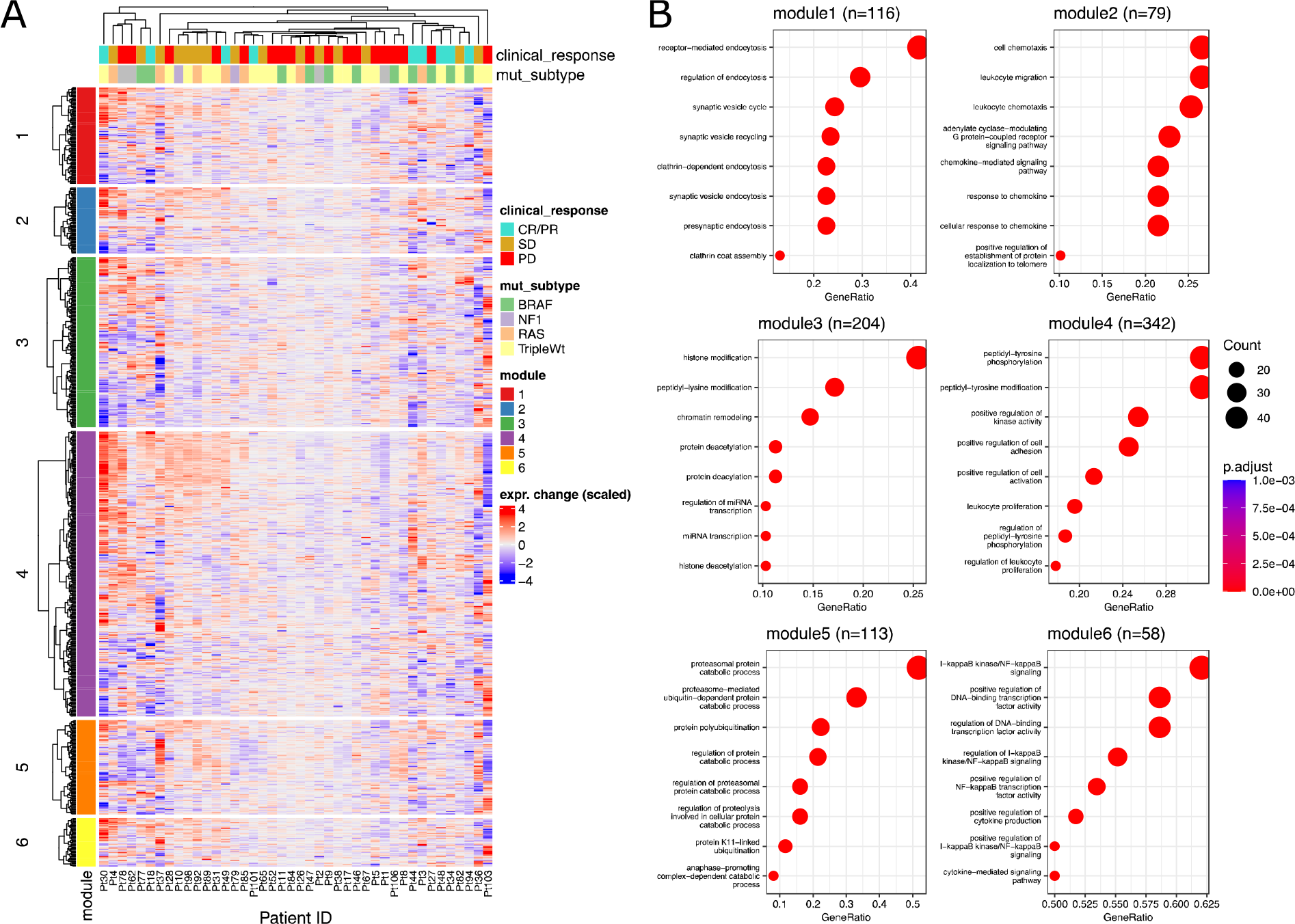
Correlated Gene Modules Enrich for Distinct Biological Functions. **A** Heatmap of scaled gene expression difference (on-treatment minus pre-treatment) for each patient, including annotation for clinical response and mutational subtype. **B:** Enrichment dotplots of top 8 most significant pathways for each gene module.

We next hypothesized that these gene modules, when differentially regulated in response to immunotherapy, may directly affect clinical outcomes. To assess the biological and clinical relevance of each module, we estimated the relative change in expression of each gene cluster in response to therapy by defining a module score as the scaled expression difference (on-treatment minus pre-treatment) averaged over all genes within each module (Fig. 3A). We assessed the relationship between these scores and patient survival by multiple Cox regression, finding that module 6 indicated a significant association with reduced risk and hence improved survival (Fig. 3B,C).

**Figure 3:**
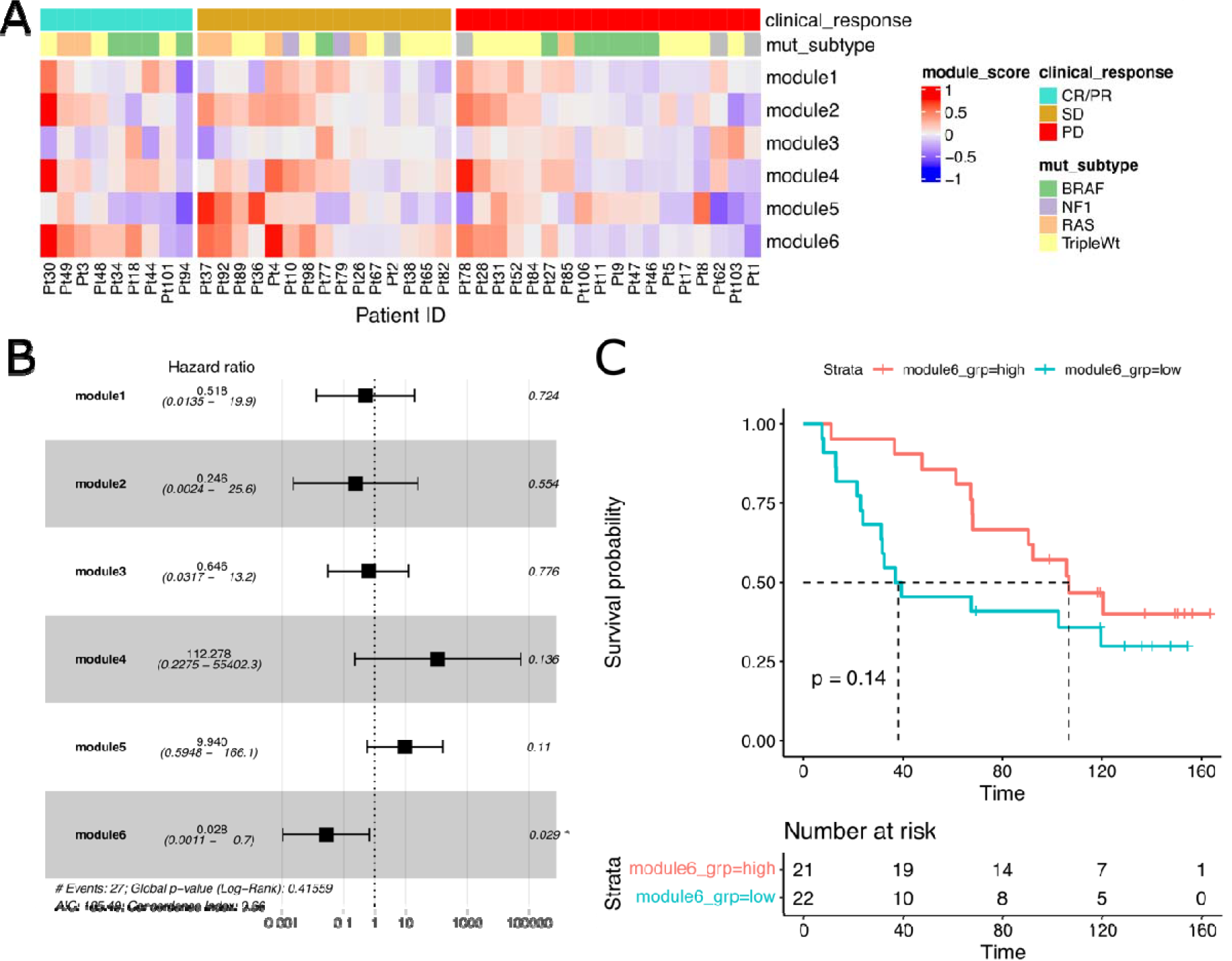
Correlated Gene Modules Associate with Survival and Clinical Response. **A:** Heatmap of module scores (as average scaled expression change of module genes) for each patient, including annotation for clinical response and mutational subtype. **B:** Forest plot of multiple Cox regression of survival with each module score. **C:** Kaplan-Meier survival curves with groups split by module 6 score median. P-value indicates log-rank test according to median high/low group. **D:** Waterfall plot of module 6 scores colored by clinical therapy response (CR/PR=complete response/partial response, SD=stable disease, PD=progressive disease).

Additionally, we observed a borderline-significant association of module 6 scores with observed clinical response to therapy (response, stable disease, or progression), whereby the module 6 score exhibited greater change on average for patients with complete or partial response and lower on average for patients with progressive disease (average module 6 score: CR/PR=0.267, SD=0.195, PD=0.019; Kruskal-Wallis test: p=0.061; Fig. 4A). We further examined the average expression change of each gene with respect to clinical treatment response, observing differential patterns for each response group wherein genes with greatest change in CR/PR responders had relatively little change in PD non-responders (Fig. 4B). Of the 58 genes in module 6, two genes exhibited significant differences (after multiple testing correction) in expression change between responders (CR/PR) and non-responders (PD); IL18R1 showed a greater positive change in responders (IL18R1 average expression change: CR/PR=0.540, PD=–0.160; FDR<0.001) while IL1RAP showed a greater negative change (IL1RAP average expression change: CR/PR=–0.781, PD=0.259; FDR<0.005). Together, these results suggest that at least one of the identified gene modules (module 6) can be associated with prognostic clinical outcomes including survival and treatment response.

**Figure 4:**
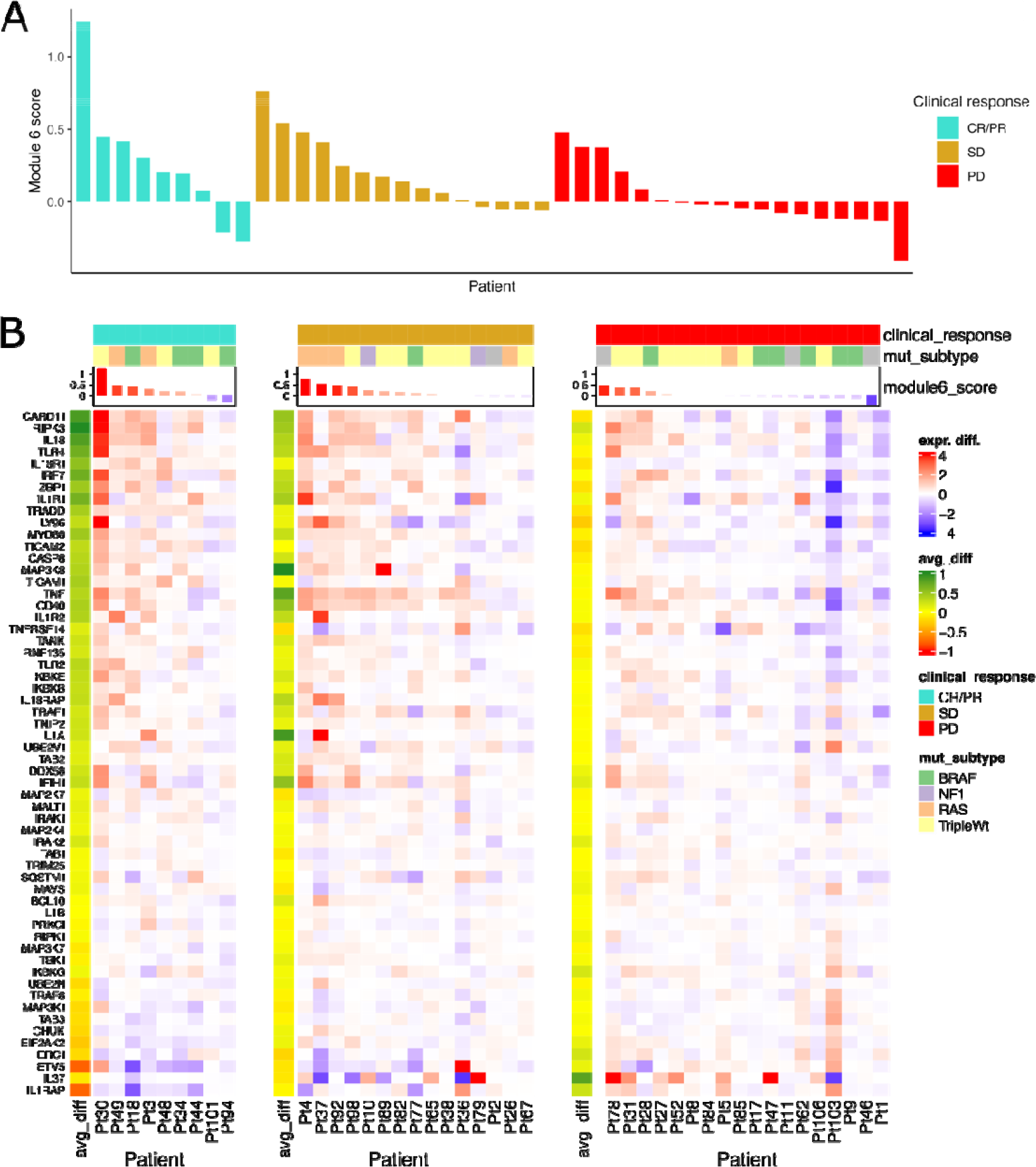
Module 6 Gene Expression Changes are Associated with Clinical Response. **A:** Waterfall plot of module 6 scores colored by clinical therapy response (CR/PR=complete response/partial response, SD=stable disease, PD=progressive disease). **B:** Heatmaps of expression change (on-minus pre-treatment) of module 6 genes in each clinical response group, including left annotation for each response group of each gene’s average expression change within the group (avg_diff).

## Discussion

In this study, we applied a network analysis approach to study correlated changes in gene expression of melanoma tumors in response to immunotherapy treatment. Using this network analysis approach, we aimed to study the complex changes that arise in genes with shared biological interactions that are dynamically regulated upon treatment induction. We considered a multi-scale geometric aspect of the gene correlation network in order to identify modules of correlated genes^18, 21^. Crucially, Ollivier-Ricci curvature (ORC), a measure that indicates how close two distributions in a network are, allowed us to distinguish within-cluster (positive curvature, more similarity) edges from between-cluster (negative curvature, less similarity) edges and accordingly classify six distinct gene modules.

The approach we applied here was similar to previous gene correlation network algorithms, including Weighted Gene Co-expression Network Analysis (WGCNA), but does not require any potentially arbitrary correlation threshold and instead directly utilizes geometric properties of the correlation network^15^. It is important to note that the Wasserstein (earth-mover’s) distance computation, which was applied to compute ORC, is effective for studying small or medium gene networks (less than about 1,000 genes) but does exhibit increasing computational time with network size. Larger gene networks on the order of several thousands or tens of thousands genes (for example, the entire set of ∼20,000 genes typically measured by RNA-seq) may become computationally burdensome or infeasible, but this might be circumvented with approximate solutions such as the entropy-regularized Sinkhorn algorithm^26^.

In terms of melanoma biology, our approach identified six distinct gene modules that represented sets of genes with shared protein interactions and correlated expression changes in response to nivolumab immunotherapy. Pathway analysis highlighted biological processes represented by the genes of each module, where we found enrichment of diverse biological processes encompassing endocytosis, chemokine signaling, histone modification, leukocyte proliferation, proteosomal catabolism, and nuclear factor kappa-B (NFkB) signaling.

We further hypothesized that these modules might be directly involved in melanoma response to immunotherapy, whereupon we identified one key module (module 6) in which a positive expression change was associated with improved patient survival and clinical treatment response. This relevant module was enriched for cytokine production and signaling but enriched even more for NFkB signaling, a pathway with known involvement in cancer immune signaling^27^. Biologically, NFkB is known as a complex of proteins which regulates inflammatory response and apoptosis in a complex manner, and thus has been implicated in cancer promoting tumorigenesis (when expressed within cancer cells) as well as anti-tumor immune response (when expressed within immune cells)^28^. Recent literature has furthermore indicated NFkB as a biomarker of clinical benefit to nivolumab in renal carcinoma^29^.

Thus, these results may be considered confirmatory of biologically relevant markers of immunotherapy and might further suggest potentially unstudied genes and mechanisms involved in melanoma immunotherapy response. In summary, we believe this study demonstrates the relevance of network curvature as a practical means of identifying gene modules in correlated biological gene expression data, and we expect this approach may be a valuable tool to study other types of cancers or other biological contexts.

## Methods

### Melanoma immunotherapy dataset

Publicly available gene expression data of 109 melanoma tumors in response to nivolumab treatment was accessed at NCBI GEO, accession code: GSE91061. Of the samples, 51 samples correspond to pre-treatment and 58 samples to on-treatment with nivolumab, with 65 patients total including 43 patients matched in both pre- and on-treatment conditions. RNA-seq data was provided both as raw gene read counts and data normalized by regularized-log normalization^30^. Entrez gene IDs were mapped to HGNC gene symbols with the org.Hs.eg.db annotation package^31^. Before downstream analysis, lowly expressed genes were removed if the gene had than less than 10 raw RNA counts in more than 90% of samples, and then rlog values of each sample were quantile-normalized to make the distribution of expression values comparable between samples.

Additional patient metadata (including therapy response and overall survival) was downloaded from the Supplemental Information of the same study^10^. Therapy responses were reported in the metadata as RECIST v1.1 categories: CR (complete response), PR (partial response), SD (stable disease), PD (progressive disease), or NE (not evaluated)^32^.

### Gene Correlation Network Construction

We constructed a weighted network model beginning with the human protein-protein interaction (PPI) network topology that represents the system of molecular interactions possible in human cells, encompassing signaling and metabolic pathways which may be modulated in various cancers. We accessed PPI topology data from STRINGdb (version 11), a database of known PPIs^25^. We incorporated a cutoff filter using the STRINGdb-provided confidence scores and a sparsification procedure based on gene ontology labels of adjacent cellular compartments to remove likely false positive edges, as previously described^13, 33^. To remove the influence of unreliable low-degree vertices, we excluded all genes with corresponding interaction degree initially less than 5. To focus our analysis, we additionally selected known immunotherapy-relevant genes involved in PD1 blockade therapies, according to the Molecular Signatures Database (MSigDB) C2 Curated Collection gene set: MsigDB C2: WP_CANCER_IMMUNOTHERAPY_BY_PD_1_BLOCKADE^22-24^. This gene set contained 23 genes (which we refer to as ‘core’ immunotherapy genes), 20 of which were contained in the expression data and STRINGdb network. We then extended the core immunotherapy gene set by including all genes with neighboring PPI edges to the core genes, for a total of 928 neighbors. Finally, we took the largest maximally connected component of the network containing these selected genes, of which 18 belonged to the core immunotherapy gene set and the remainder were PPI neighbors. These criteria resulted in an undirected network topology with 912 vertices (genes) and 50,518 edges.

Using the difference of (rlog) normalized gene expression in on-treatment minus pre-treatment, a weighted correlation matrix *C* was computed representing the Pearson correlation of all patients’ expression change for each pair of genes, then shifted from the range [-1,1] to the range [0,1] by a linear transformation 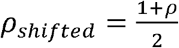, as a similarity metric such that negative correlations become close to zero and positive correlations remain close to 1. To define transport cost on the network (as utilized below in the Wasserstein computation), correlations were transformed into distance-like edge weights defined as the inverse of the shifted correlation if that edge was in the given PPI network topology. Then, a distance matrix *d* representing shortest path length between each pair of genes was computed using Djikstra’s algorithm^34^.

### Dynamic Network Curvature Analysis

Dynamic network curvature analysis was conducted by simulating diffusion over the weighted network and measuring geometric changes^18, 21^. First, the graph Laplacian *L* was determined as

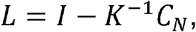

where *I* is the identity matrix, *C*_*N*_ is the shifted correlation matrix of edges in the network and *K* is the weighted degree or row-sum of *C*_*N*_.

The graph Laplacian represents the divergence of weighted differences in a discrete graph and served as a crucial tool to efficiently simulate diffusion at multiple pseudotime/scale parameters *τ*. A diffusion distribution matrix *D* was computed using the matrix exponential of *L*:

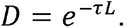

Each row of *D* indicates a probability distribution corresponding to one diffusion process with an initial Dirac delta *δ* _*i*_ concentrated at a single vertex *i* then diffused over pseudotime *τ* to arrive at a diffused distribution. We applied this step for 101 values of *τ* ranging in the form log_10_(*τ*) ∈[-2,2].

In each diffused graph, Ollivier-Ricci curvature (ORC) *κ* _*i*_ was computed for each edge in the graph by first computing the Wasserstein distance *W*_1_ between two probability distributions:

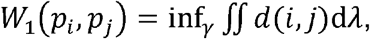

as the minimum total cost for all couplings *γ* that satisfy marginals *p*_*i*_ and *p* _*j*_ signifying probability distributions of vertex *i* and *j* diffused for the same pseudotime *τ* and transport cost *d* specified by the shortest path length between each vertex. The Wasserstein distance (*W*_1_) thus indicates optimal transport distance as a measure of closeness between two distributions. Then, ORC subsequently transforms this value by the following formula:

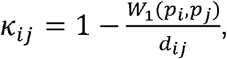

where *d*_*ij*_ is the direct distance between the two vertices defined above. Curvature *κ* can indicate positive convergence (clique-like) or negative divergence (tree-like) of the probability distributions in the graph, revealing the geometric structures of the graph (i.e. clusters, branching).

Over the diffusion process, *κ* initially begins at zero, indicating the transport distance of concentrated deltas at each vertex is equivalent to the direct edge distance between them. As the diffusion process progresses to fully diffused stationary distributions, there will be little to no transport distance as the distributions become equal, so *κ* will approach 1. It is in the middle of the diffusion process, however, that *κ* can become negative for certain edges or remain positive for others, thereby revealing critical network edges that point to overarching community structure within the graph. After measuring all edge curvature values over the diffusion evolution, we determined a threshold *τ*_*crit*_as the first pseudotime/scale when the upper 99^th^ percentile of all edges exceeded *κ* ≥ 0.75. For each edge, we determine *κ*_*crit*_ to be the value of *κ* at *τ*_*crit*_. We additionally define 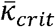as the integral 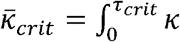, to represent a smoothed estimate of curvature during diffusion up to *τ*_*crit*_.

### Gene Module Clustering

At the critical threshold *τ*_*crit*_, all edge 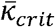 values were considered as modularity weights in a weighted Louvain clustering algorithm, such that the Louvain optimization iteratively maximized 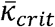 within clusters and minimized 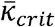 between clusters^35^. This was accomplished using the networkx Python package implementation of nx.community.louvain_communities, with the default resolution parameter of 1, which ultimately assigned each gene an integer label corresponding to one of six gene modules^36^.

With each module, a module score was computed for each patient that could be considered to assess how each module score related to clinical characteristics including overall survival and therapy response. Scaled gene expression difference was defined as the difference in normalized gene expression (on-treatment minus pre-treatment condition in each patient with matched data for both conditions), then scaling each gene by dividing by standard deviation across all patients (but not shifting the mean, as typical for z-score, so as to the preserve positive and negative sign of expression change). Gene module scores were then computed for each patient as the average scaled gene expression difference over all genes within each module. Given the biological context of the correlation network in melanoma immunotherapy response, we applied pathway analysis on each gene module to identify biological processes involved in each module. Gene ontology (GO) enrichment was computed using the clusterProfiler R package (considering “ALL” pathways of GO BP, CC, and MF subontologies) and the enrichplot R package was utilized for visualization of pathway analysis results^37, 38^.

### Statistical Analysis

An unpaired t-test was used to compare curvature of within-cluster vs between-cluster edges. Pathway enrichment analysis utilized a hypergeometric test for over-representation analysis, including multiple hypothesis correction, for which we select a cutoff of FDR>0.05^39, 40^. Multiple Cox proportional hazard test was applied to determine the association of all module scores with patient survival. Kaplan-Meier analysis of module 6 score split into low/high groups by median was used for visualization. For statistical analysis related to therapy response, we removed 1 patient with NE and grouped CR (n=3 patients) and PR (n=6 patients) as a single group CR/PR. Kruskal-Wallis test was used as a non-parametric analysis of variance to assess module score association with therapy response. For each of 58 genes in module 6, a two-sample Wilcoxon test was applied to compare expression change in CR/PR vs PD groups, followed by a significance cutoff of FDR<0.05 after Benjamini-Hochberg multiple-comparison adjustment^40^.

## Code Availability

All analysis code was written in Python and R and has been made publicly available in a GitHub repository at the following link: <https://github.com/kevin-murgas/melanoma_dynamic_curvature>.

## Acknowledgements

Research partially supported by the Breast Cancer Research Foundation Grant BCRF-17-193 (K.A.M., R.E., J.O.D., A.R.T.), AFOSR FA9550-20-1-0029 and FA9550-21-1-0096 (K.A.M., R.E., A.R.T.), ARO W911NF2210292 (K.A.M., R.E., A.R.T.), Memorial Sloan Kettering Cancer Center Support grant P30-CA008748 (K.A.M., R.E., J.O.D., A.R.T.), and GIF Research Grant No. I-1514-304.6/2019 (E.S.).

